# Identifying the Fusarium species involved in foot rot disease of faba beans in the UK using a combined molecular and microbiological approach

**DOI:** 10.1101/2024.10.25.620281

**Authors:** Basem Attar, James J. N. Kitson, Jordan P. Cuff, Becky Howard, Ana Lages, Dina Gomez, Neil Boonham

## Abstract

Foot rot is a devastating disease of faba bean crops globally, including in the United Kingdom, the world’s third largest producer. To identify the causal agents, we have sampled foot rot-affected plants and soils from faba bean crops across England. We isolated organisms associated with foot rot disease in culture and assessed pathogenicity *in vivo* to evaluate the infectivity of the isolates on faba bean. We identified the pathogenic isolates using DNA barcoding of the Internal Transcribed Spacer (*ITS*) and Translation Elongation Factor one α (*TEF1* α) molecular markers. A total of 113 clonal isolates were obtained from infected plants and soil samples across England. Of these, 60 were pathogenic, inducing mild to severe symptoms on faba bean. Sequencing of the ITS and *TEF1* α loci and comparison against sequence databases (Genbank and Fusarium_ID) enabled the identification of pathogenic isolates, in decreasing order of frequency, as *Fusarium oxysporum* (26.6 %), *F. vanettenii* (25%), *F. redolens* (15 %), *F. solani* (11.6%), *F. culmorum* (8.3 %), *F. avenaceum* (6.7 %), *F. equiseti* (1.7 %), *F. clavum* (1.7 %), *Clonostachys rosea* (1.7%) and *Alternaria alternata* (1.7%). *F. oxysporum, F. redolens* and *F. avenaceum* induced the most severe symptoms, whilst *F. solani* induced the least severe symptoms. Determining the most prevalent causal agents of foot rot in UK faba beans will facilitate targeted disease monitoring and intervention for enhanced productivity.

## Introduction

Faba bean is one of the most important legume crops globally, grown as a source of plant-based protein and an essential component of successful crop rotation (Emeran et al., 2011; de Visser et al., 2014). According to the FAO, the United Kingdom was the third largest global producer and second largest exporter of faba bean during the 2019 planting season, responsible for 547,800 and 119,071 metric tons, respectively (FAO 2020). Faba bean is, however, affected by many diseases and pests that limit its global production. The most impactful diseases are chocolate spot, caused by *Botrytis fabae*, Ascochyta blight, caused by *Ascochyta fabae*, rust, caused by *Uromyces vicia fabae*, and the foot and root rot disease complex, the causal agents of which are poorly characterised (Sillero et al., 2010a).

Root rot diseases are important in a broad spectrum of crops worldwide and can be caused by a range of pathogenic organisms, including fungi, bacteria, viruses and oomycetes (Gonzalez et al., 2011). The main symptoms are brown to black spots on the crown of the plant and the base of the stem, black spots on the top of the root, and absence of rootlets. The affected plants appear yellow with gradual wilting of the upper tissues. The symptoms become more severe with time, eventually killing the plant (Gonzalez et al., 2011; Nzungize et al., 2011).

Foot and root rot on faba bean is reported as a limiting factor for production in many areas (Infantino et al., 2006), including Canada (Chang et al., 2014), Ethiopia (Bogale et al., 2009) and China, where the disease is reported to cause 100 % yield loss when climate conditions are suitable for disease development (Zhang, Li & Guo, 2018). Many organisms have been associated with this disease complex, including members of the genus *Fusarium*, which are frequently isolated from infected faba bean plants (Figure 1). These include *F. oxysporum* Schlecht, *F. solani* (Mart.) Sacc., *F. avenacearum* (Corda ex Fr.) Sacc., *F. graminearum* Schwabe and *F. culmorum* (Smith) Sacc. (Helsper *et al*., 1994; Sillero *et al*., 2010a), but comprehensive surveys are lacking in most countries, including the UK, and other species are likely to be involved. In addition to *Fusarium*, organisms from many other genera have been reported to be involved in foot and root rot, including *Rhizoctonia solani* Kuehn, *Pythium* spp., *Phoma* spp., and *Aphanomyces eutichus* Drech. (Salt, 1982; Sillero *et al*., 2010b).

**Figure 1:**
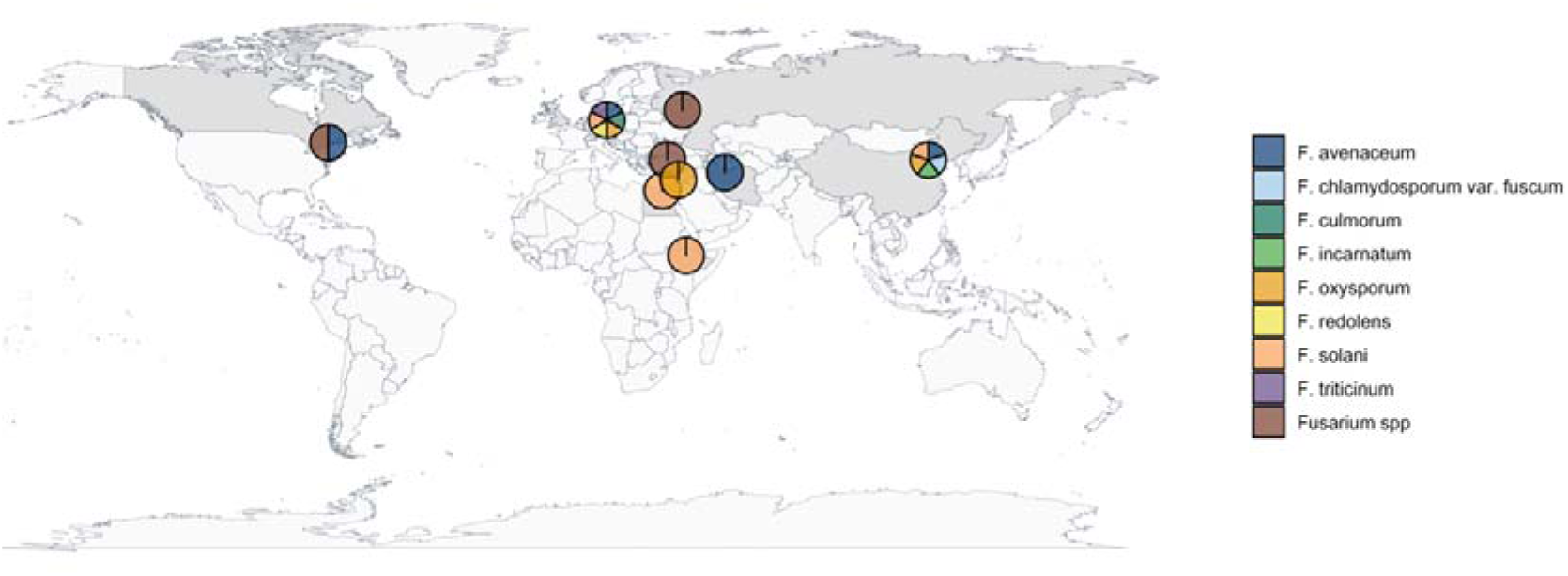
The Fusarium species recorded as causal agents for foot and root rot disease complex globally.

Given the involvement of several species in foot rot, understanding the factors that influence which species predominate is important for guiding monitoring and management. Differences in climate conditions can prioritise the abundance of some species over others; for example, in China, many species were reported as causal agents for foot and root rot, but *F. solani* was more frequent in Qinghai, *F. acuminatum*, *F. oxysporum* and *F. moniliforme* were more frequent in Zhejiang, and *F. oxysporum*, *F. avenaceum*, *F. moniliforme* and *F. equiseti* were more frequent in Jiangsu (Ren, 2003; Rubiales & Khazaei, 2022; Yu et al., 2023). This indicates that region- and context-specific surveys are required to understand the likely causal agents for optimal monitoring and management. Although the foot and root rot disease complex is established in the United Kingdom (Clarkson, 1978), the causal agents are poorly characterised, and more granularity of surveys and information are required to design effective disease monitoring and management strategies in the future. Here, we identify the dominant causal agents of faba bean foot and root rot disease in England using a combination of microbiological and molecular approaches. We specifically aimed to characterise the species responsible for foot and root rot, the frequency of their involvement across a nationwide survey and the disease severity that each species caused. This information will be invaluable for guiding future monitoring and management schemes to reduce the impact of foot and root rot for UK faba bean production.

## Materials and methods

### Fungal isolation

Isolates (113) were prepared from both soil and infected plant samples that were received from the Plant Clinic at the Processors and Growers Research Organisation (PGRO). The samples were from different regions of England, United Kingdom (Table 1).

**Table 1:** The locations and number of isolates obtained for infected faba bean and soil samples used in the study. * = includes soil isolates.

| Location | Number of isolates obtained | Month(s) |
| --- | --- | --- |
| PGRO experimental plots | 29* (13 soil, 16 plant) | November 2022 |
| Cambridgeshire | 2 | June 2023 |
| Oxfordshire | 3 | June-July 2023 |
| Durham | 3 | July 2023 |
| Shropshire | 4 | July and August 2023 |
| Lincolnshire | 5 | July and August 2023 |
| Northumberland | 5 | July and August 2023 |
| Staffordshire | 4 | July and August 2023 |
| Suffolk | 4 | July 2023 |
| Leicestershire | 2 | July 2023 |
| Essex | 4 | July 2023 |
| Norfolk | 34 | August 2023 |
| Yorkshire | 6 | September 2023 |
| Hampshire | 2 | July 2023 |
| Undisclosed PGRO locations | 6 | Undisclosed |

Infected plant samples were disinfected by placing pieces of infected stems/root in 10 % sodium hypochlorite solution for 5 min. The samples were rinsed twice using sterilised distilled water and placed on sterilised filter paper to be dried for 10 min at room temperature inside a laminar flow hood. The dried samples were moved to potato dextrose agar medium (PDA) inside 9 cm diameter plastic Petri dishes. Petri dishes were incubated at 22 °C with 12 h fluorescent photoperiod, and light intensity of 20-25 µMol.m^-2^.s^-1^.

Once colonies had formed, clonal isolates were prepared from the colonies as follows. Approximately 1 mm^2^ of the colony was collected using a flame-sterilised inoculation loop and was sequentially spread onto three Petri dishes containing 2 % water agar, to dilute the inoculum gradually. The Petri dishes were incubated for 24 h as described above. The third Petri dish for each isolate was examined under a stereoscope, and one separated hypha was transferred using a flame-sterilised scalpel to another Petri dish containing PDA medium, and incubated for seven days to provide a clonal isolate. The isolates were stored for future use using two methods, for routine or long-term storage. For routine storage (months), three discs of the PDA medium containing the clonal isolate were transferred to a 2 ml microcentrifuge tube containing 1 ml of sterilised distilled water, the tube sealed with parafilm and stored at −20 °C. For long-term storage (3-4 years) the clonal isolate was plated onto a Petri dish containing many pieces of 1 cm long sterilised filter paper on PDA medium, and the colony was allowed to grow for seven days to cover the filter paper. The pieces of filter paper were removed and placed inside an empty Petri dish and dried for seven days at room temperature. The filter paper pieces were then transferred to an empty 2 ml plastic microcentrifuge tube and stored at −20 °C. Koch’s postulates were confirmed for each of the isolates by re-isolation, inoculation, and identification.

### Pathogenicity testing

Pathogenicity testing was conducted using susceptible faba bean seedlings (cv. Lynx) grown in test tubes in a mixture of perlite/vermiculite. The growth media was prepared by adding one volume of vermiculite (the capacity of a 1000 ml plastic beaker) to one volume of perlite inside an autoclave bag; this was mixed to ensure equal distribution of each component, and 1 litre of distilled water was added. The autoclave bag was closed and autoclaved for 20 min at 121 °C, and the mixture was transferred to fill 2/3 of the test tubes (150 x 24 mm, 1.2 ml wall; borosilicate glass 150 x 24 mm, rimless, Appleton Woods Ltd), which were then sealed with cotton wool and aluminium foil, prior to being autoclaved.

Seeds were soaked in sterilized distilled water overnight and placed in 10 % sodium hypochlorite for 5 min. The seeds were washed three times with sterilised distilled water and placed on sterilised filter paper until dry. The seeds were then transferred to 9 cm petri dishes containing 1.2 % Tap Water Agar (12 g agar in 1 l of tap water, autoclaved in a 2 l conical flask), where they were allowed to germinate for four days in the incubator at 24 °C before being transferred to test tubes containing the vermiculite/perlite mixture.

Following transfer, the seedlings were allowed to grow for five to seven days until they were suitable for inoculation (4-5 cm root length). The seedlings were inoculated by placing a 10 mm block of PDA medium containing a ten-day old fungal culture against the stem base. A small piece of sterilised cotton was placed around the stem to ensure adequate moisture at the inoculation site. The inoculated seedlings were incubated at 24 °C, with a 12 h photoperiod and light intensity of 20-25 µMol.m^-2^.s^-1^.

Disease severity was monitored daily from 5-6 days after inoculation. Root and stem infection were scored on days 15 and 25 using a 5-point scale (Figure 2):

**Figure 2:**
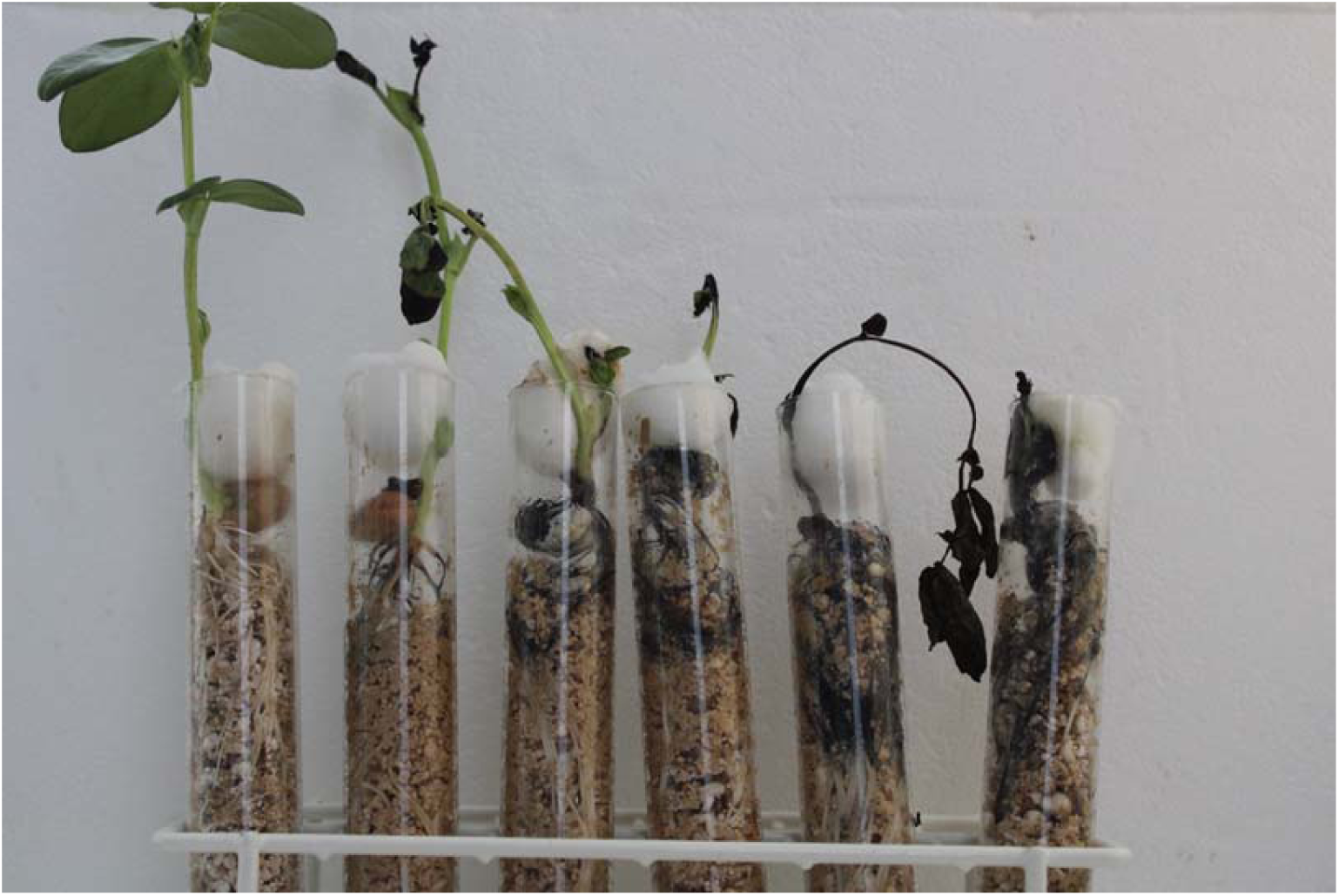
From left to right, the 0-5 scale that was used to score the pathogenicity and aggressiveness of the fungal isolates in in vitro experiments.

0 = healthy roots, no discolouration.

1 = up to 20 % root or stem base discoloured.

2 = 20-40 % root or stem base discoloured.

3 = 40-60 % root or stem base discoloured.

4 = 60-80 % root or stem base discoloured, stunting of plant.

5 = total discoloration, dead plant.

### DNA extraction

Clonal colonies were cultured on 50 ml of Potato Dextrose Broth (PDB) medium (FORMEDIUM^TM^) in a 250 ml flask and incubated for seven days on a rotary shaker (22 °C with 70 RPM). The mycelium for each isolate was harvested by transferring the contents of each flask into a 50 ml falcon tube and centrifuging at 5000 RPM for 10 min. The supernatant was discarded, and the mycelium pellet stored at −20 °C. Mycelium (1 ml) was transferred to a 2 ml safe-lock microcentrifuge tube. The tubes were covered with parafilm which was pierced to allow moisture to evaporate. Samples were freeze-dried (−45 °C, 0.133 mbar) for 48 h. Freeze-dried mycelium (15 mg) was transferred to a 2 ml safe lock microcentrifuge tube containing two carbon steel ball bearings (3 mm diameter, grade 1000, SimplyBearings). The mycelium was homogenised for 4 min using a TissueLyser (Retsch MM400; 30 RPS), after which 120 µl of TNES buffer was added and the samples were homogenised again as before.

Total DNA was extracted from the samples following the BOMB-Bio nucleic acid tissue DNA extraction protocol (Oberacker et al., 2019). The samples were incubated at 55 °C overnight after adding 2 µl of proteinase K and 3 µl of RNAase A. Following incubation, 240 µl of GITC lysis buffer was added, mixed and samples incubated at room temperature for 5 min. Isopropanol (480 µl) was added and, following centrifugation (5000 RPM; 5 min), 650 µl was transferred to a new tube with 200 µl of 1X BOMB-Bio magnetic bead solution (1:50 carboxylated SeraMag Speed Beads magnetic beads in TE) and mixed. The solution was placed on a magnetic rack to hold the beads with DNA bound to them in place whilst the supernatant was removed. The beads with bound DNA were washed once with isopropanol (400 µl) and twice with 80 % ethanol (400 µl). The solution was removed from the magnet, beads were allowed to dry briefly and 70 µl of nuclease free water was added to elute the DNA. The beads were removed by placing the samples back on the magnetic rack and transferring the supernatant to a new tube. The extracted DNA concentration was estimated using a Qubit 4 and 1X dsDNA High Sensitivity (HS) Assay Kit (Invitrogen) according to the manufacturer’s instructions.

### Polymerase Chain Reaction (PCR)

DNA was amplified using PCR with three sets of primers: Internal Transcribed Spacer (ITS) primers ITS1/ITS4 (Raja et al., 2017) and two sets of Translation Elongation Factor one α primers, 1018F/1620R and EF1/EF2 (O’Donnell et al., 1998; Raja et al., 2017). Prior to PCR, the template DNA concentration was adjusted to 5 ng/μl using molecular grade water. The PCR mix contained: 12.5 µl of 2x MyTaq Red Mix (Meridian Bioscience), 3 µl of template DNA, 1 µl of each primer (10 µM) and 7.5 µl of nuclease free water to a final volume of 25 µl. The cycling conditions and sequence of each primer are given in Table 2. The PCR products were separated alongside a 1 kbp ladder (GeneRuler 1 kb Plus DNA Ladder Thermo scientific SM1331) using gel electrophoresis in a 1 % agarose gel in TAE buffer stained with GelRed® nucleic acid stain (Sigma Aldrich). Gels were visualised with a UV transilluminator (BioRad) to confirm successful amplification.

**Table 2:**
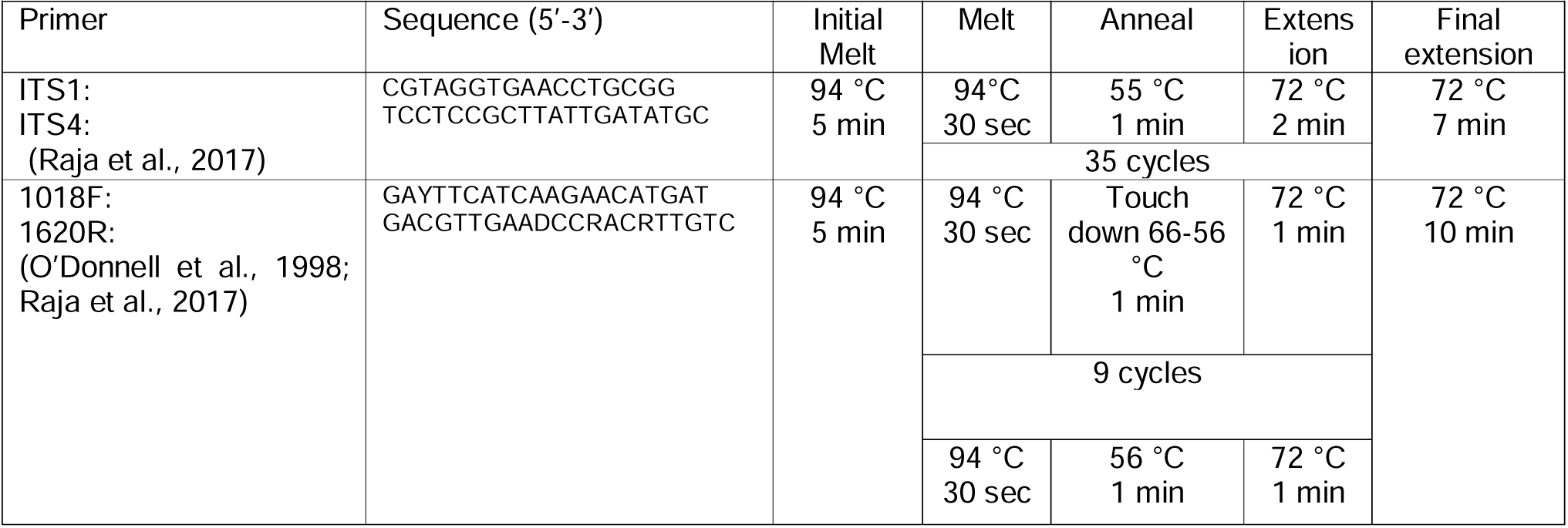
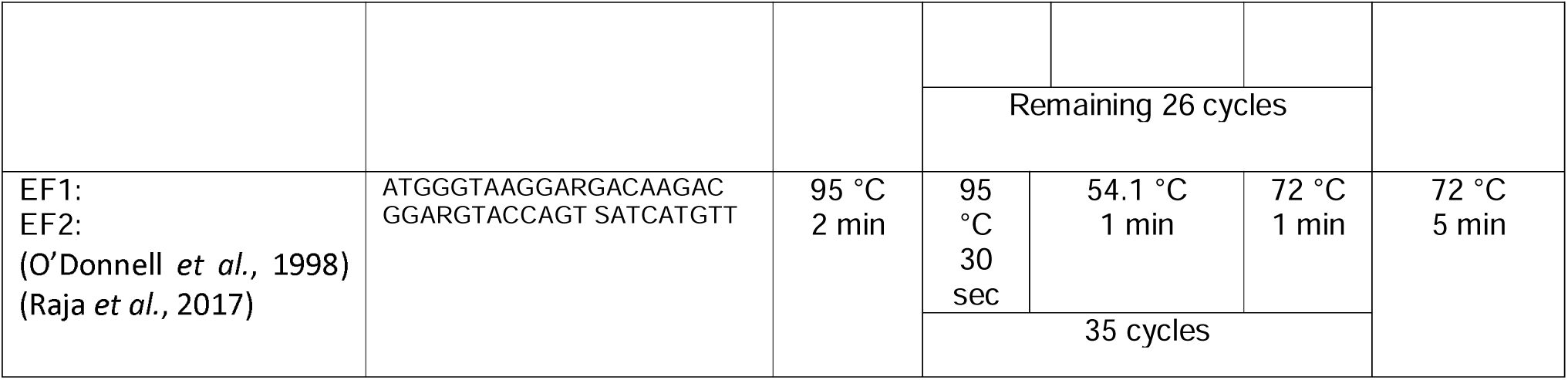
The ITS (ITS1 and ITS4) and TEF1α (1018F, 1620R, EF1 and EF2) primer sequences and PCR conditions.

Prior to sequencing, the PCR products were purified using solid-phase reversible immobilisation (SPRI) magnetic beads. To each PCR product, 1 X SPRI beads were added at a 1:2 ratio of PCR product to bead solution (50 µl PCR product to 100 µl SPRI beads) and the mixture incubated for 5 min at room temperature. The tubes were transferred to the magnetic plate for 5 min until the solution was clear, at which point the supernatant was discarded and the beads were washed twice with 80% ethanol for 60 s each time. Ethanol was removed and the beads held on the magnetic plate were allowed to dry for 5-10 min at room temperature. The tubes containing the dry beads were removed from the magnetic plate, and 50 µl of molecular grade nuclease free water was added to each tube, mixed and incubated for 5 min, allowing the DNA to elute. The tubes were placed back on the magnetic plate for 5 min until the solution was clear, at which point the supernatant was transferred to a new 1.5 ml microcentrifuge tube. Sanger sequencing was carried out using ITS1, ITS4 primers and EF1, EF2 primers. All sequencing was carried out by Eurofins Genomics.

Following sequencing, the chromatograms obtained were trimmed and analysed using Geneious Prime (version 2023.0.4). First, low-quality bases (e.g., overlapping peaks) were trimmed from each end and the sequence upstream from that site, including the primer sequence, was deleted. The forward and reverse sequence for each sample were assembled using the *de novo* assemble function in Geneious at the highest sensitivity to create a consensus sequence (highest threshold quality: 60%). A basic local alignment search tool (BLAST) search was carried out for the consensus sequences using the NCBI-NR database for the ITS sequences, and, for the TEF1α sequences, the data available on the Fusarium ID database (Torres-Cruz et al., 2022). All of the *TEF1*α consensus sequences from all samples were subsequently aligned with the available sequences on the Fusarium ID database (Geneious global alignment with free end gaps, 65% similarity), and a phylogenetic tree was generated (genetic distance model Tamura-Nei, neighbour joining method, bootstrap with 100 replicates).

## Results

### Visual inspection, microscopy and identification of pathogenic isolates

The visual inspection of colonies on PDA using microscopy highlighted that different *Fusarium* colonies differed in colour and the density of micro- and macro-conidiospores. The colonies appeared lighter (a pinkish white colour) with fluffy (cotton-like) aerial mycelium for *F. avenacium*, *F. solani* and *F. culmorum* isolates, pinkish white for *F. oxysporum* and *F. redolens* isolates, and dark green with less obvious aerial mycelium for *F. vanetenii* isolates. The pathogenicity tests for the 113 isolates showed that 60 were pathogenic on the susceptible faba bean, with a range of disease severities, whilst 53 did not produce disease symptoms, the identities for which are given below. Koch’s postulates were confirmed for each of the isolates, by re-isolation, inoculation and identification.

### DNA barcoding results and phylogenetic analysis

The ITS and *TEF1*α PCRs produced amplicons between 600-700 base pairs and 800-900 base pairs, respectively. Following BLAST searching of GenBank using the ITS sequences, 58/60 pathogenic isolates were identified as members of the genus *Fusarium* and 2/60 were identified as *Clonostachys rosaea* and *Alternaria alternata*. The remaining non-pathogenic isolates (34/113) were identified as *Botrytis cinerea*, *Alternaria spp*, *Alternaria infectoria*, *Chaetomium* sp, *Diaporthe columnaris*, *Trichoderma atroviride*, *Trichoderma* sp, *Penicillium* sp, non-cultured fungus, *Absidia glauca*, *Chaetomium cochliodes*, *Sordaria fimicola, Didymella pinodella, Globisporangium intermedium*, *Glonium pusillum*, *Clonostachys rosea* and *Epicoccum layuense.* A further 19 isolates that had a fungal like growth *in vitro* could not be identified using either the *ITS* nor *TEF*1α loci.

To identify the *Fusarium* isolates to species, consensus sequences, trimmed to represent the same fragment of the *TEF1*α gene, were compared against the Fusarium_ID database using BLAST. The results identified the presence of eight different *Fusarium* species as follows, in order of frequnecy: *Fusarium oxysporum* (26.6 %), *F. vanettenii* (25%), *F. redolens* (15%), *F. solani* (11.6 %), *F. culmorum* (8.3 %), *F. avenaceum* (6.7 %), *F. equiseti* (1.7 %) and *F. clavum* (1.7 %; Figure 3).

**Figure 3:**
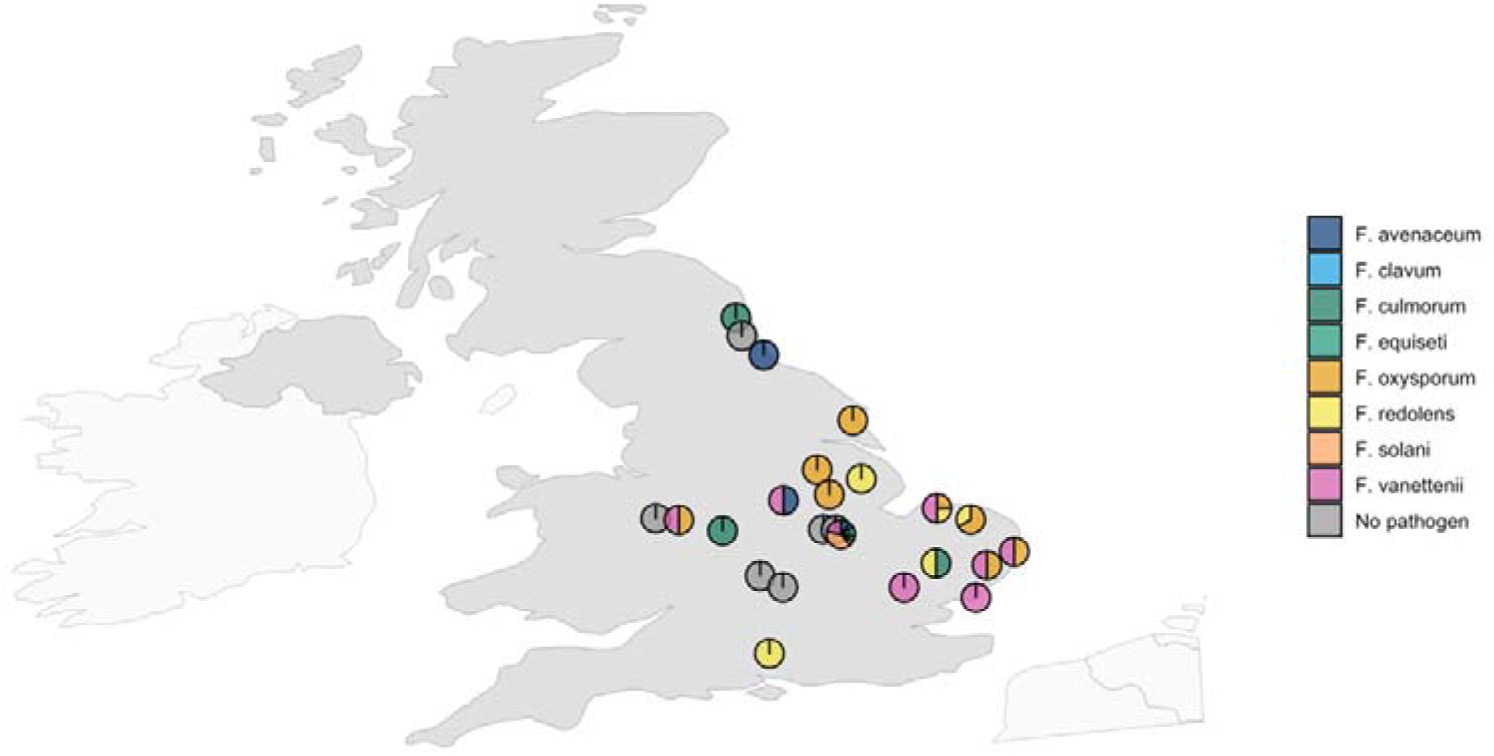
The pathogenic fusarium species that were found in England and identified by DNA barcoding and comparison to Fusarium_ID database entries

The phylogenetic analysis of the consensus sequences for the 58 isolates detected as *Fusarium,* alongside the sequences available on the Fusarium_ID database, showed that each species detected in this study was found within the same clade as conspecifics in the database (Figure 4).

**Figure 4:**
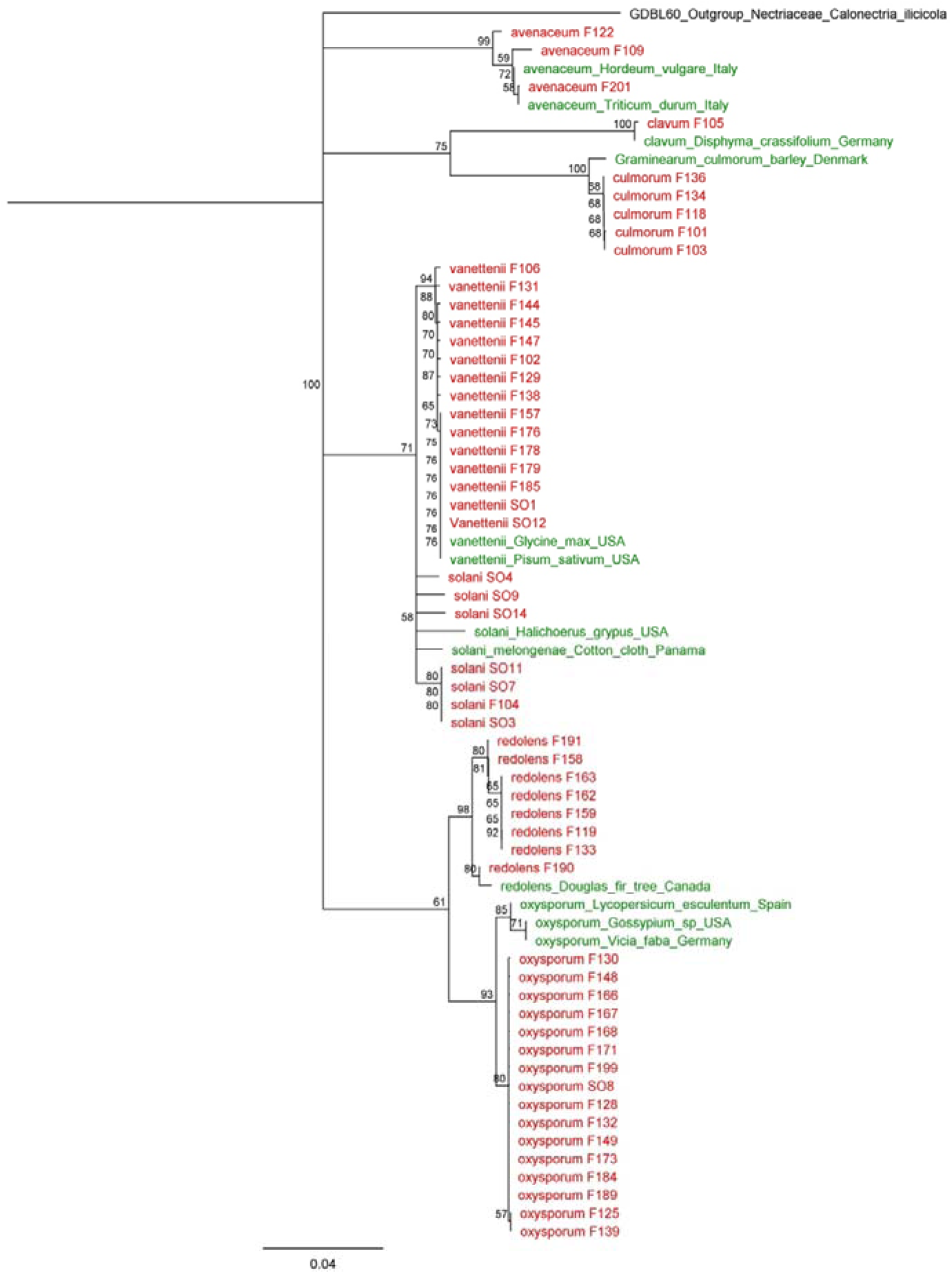
Phylogenetic tree for the isolates included in this study. The tree was generated for the sequences obtained in this study alongside sequences available in the Fusarium_ID database. Isolates used in this study are in red, whereas those from the Fusarium_ID database are given in green with their country of origin given at the end of their name.

### Pathogenicity variation

Disease severity significantly differed between *Fusarium* species (ANOVA: F = 4.200, df =7, p = 0.001). Isolates of *F. oxysporum*, *F. avanaceum* and *F. redolens* induced the most severe symptoms. The disease severity of *Fusarium solani* significantly differed from *F. oxysporum* in pairwise post-hoc tests (Tukey: difference = −2.232 ± 1.780, adjusted p = 0.005; Figure 5), *F. vanetenii* significantly differed from *F. oxysporum*, (Tukey: difference = −1.575 ± 1.411, adjusted p = 0.019; Figure 5), and *F. solani* marginally insignificantly differed from *F. redolens* (Tukey: difference = −1.968 ± 1.979, adjusted p = 0.052; Figure 5). In addition to *Fusarium*, two other species, *Clonostachys rosaea* and *Alternaria alternata,* were pathogenic on faba bean giving mild symptoms.

**Figure 5:**
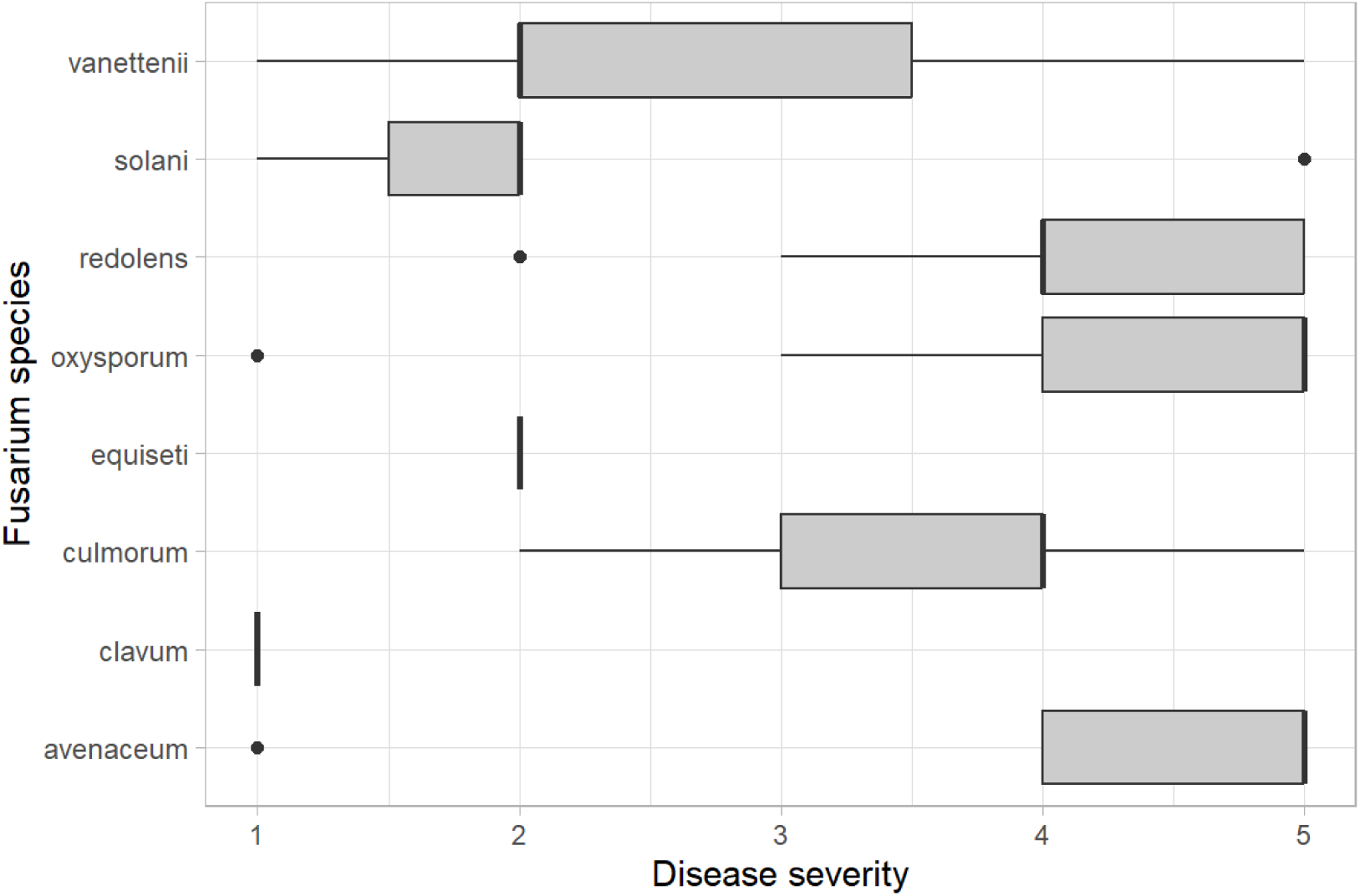
Disease severity for different Fusarium species inoculated on faba bean seedlings inside test tubes. The severity was measured on a 0-5 scale based on the extent of tissue spotting and wilting.

## Discussion

The faba bean foot and root rot disease complex detrimentally impacts faba bean production worldwide, including in the UK. Despite a geographically variable range of organisms having been identified as causal agents of the disease, this varies internationally, and, to our knowledge, our study is the first to identify the causal agents in England.

Following *in vitro* pathogenicity testing of 113 isolates from faba bean samples exhibiting typical disease symptoms collected in a nationwide survey across England, eight *Fusarium* species (*Fusarium oxysporum*, *F. vanettenii*, *F. redolens*, *F. solani*, *F. culmorum*, *F. avenaceum*, *F. equisiti*, and *F. clavum*) were identified that induced foot rot symptoms. We recovered the identified *Fusarium* species at different frequencies, with *F. oxysporum*, *F. vanettenii*, and *F. redolens* being the most frequently associated with disease symptoms. We also found significant variation in the severity of symptoms initiated by different *Fusarium* species, where *F. oxysporum*, *F. avenaceum* and *F. redolense* induced significantly more severe symptoms than isolates of other species.

The association of different *Fusarium* species with foot and root rot disease in faba bean has been reported in other global regions previously. Isolates of *F. culmorum*, *F. oxysporum*, *F. avenaceum, F. redolens* and *F. equiseti* have been reported as causal agents for the disease (Helsper et al., 1994; Sillero et al., 2010b; Šišić et al., 2022). *Fusarium* has been reported on faba bean from many countries (Figure 1), including *Fusarium* spp, and *F. avenaceum* in Canada (Lamari & Bernier, 1985; Chang, K. F., Conner, R. L., McLaren, D. L., Hwang, S. F., & Strelkov, 2010), *F. redolens*, *F. avenaceum*, *F. triticinum*, *F. culmorum*, *F. oxysporum* and *F.solani* in Germany (Baćanović-Šišić et al., 2018), *Fusarium* spp in Turkey (Erper et al., 2008), *F. oxysporum* in Syria (Akem, C. and Bellar, 1999), *F. solani* in Egypt and Ethiopia (Abou-Zeid et al., 1997; Tamiru & Muleta, 2018), *F. avenaceum* in Iran (Hemmati, 2014), *F. solani*, *F. avenaceum*, *F. oxysporum, F. incarnatum* and *F. chlamydosporum* var. *fuscum* in China (Zhang, Y., L. Li, Y. P. Guo, 2018; Chu et al, 2019), and *Fusarium* spp in Russia (Ashmarina, L. F. Gorobey, I. M. Konyaeva, N. M., And Agarkova, 2010). There was significant overlap between the *Fusarium* species associated with the disease in England and other parts of the world; however, we identified two species, *F. clavum* and *F. vanettenii,* which, to the best of our knowledge, have not previously been reported as associated with faba bean foot and root rot disease. We also identified an isolate of *C. rosaea* which was pathogenic on faba bean, confirming the findings of Afshari & Hemmati (2017). This is particularly significant as isolates of this species have been used as a biocontrol agent for the control of soil-borne pathogens (Krauss & Soberanis, 2001; Samsudin et al., 2017), suggesting that *C. rosaea* is an opportunistic fungal pathogen. This potential opportunism is evident in our study since only two of four *C. rosaea* isolates were pathogenic. More research is required to investigate the pathogenicity of *C. rosaea* isolates to identify determinants of its potential pathogenic activity. Such work should especially focus on *C. rosaea* isolates recovered from commercial biocontrol products, as the potential pathogenicity of these products could be of great concern to growers if these products pose any risk to crops.

The variable prevalence and pathogenicity of different causal agents of foot and root rot disease indicates that monitoring and management of the disease needs to be adapted for different locations. Active monitoring is required to identify these causal agents within specific locations across time. It is also important to widen surveys across the entire UK to determine whether the prevalence of specific causal agents is regionally variable, and whether this relates to local environmental conditions, or differences in management practices or faba bean varieties. While *Fusarium* is the primary causal agent of foot and root rot in many parts of the world, the frequency and importance of each *Fusarium* species is thought to be affected by local agronomic differences (Chatterton et al., 2019; Williamson-Benavides and Dhingra, 2021). In Germany, the frequency of *F. redolens* is higher in organic compared to conventionally managed fields; however, the frequency of *F. oxysporum* and *F. solani* was not affected by either location or agricultural practice (Šišić et al., 2022). In China, the frequencies of different *Fusarium* species differed between different geographic locations, with *F. avenaceum* and *F. oxysporum* being dominant in Gansu (Zhang, et al., 2018b), while *F. incarnatum* and *F. chlamydosporum* var. *fuscum* were dominant in Xinjiang (Chu et al, 2019). Further research is required to investigate links between agronomy, location and the identities of foot and root rot causal agents both in the UK and globally, which could refine predictions and management of disease outbreaks.

We found variation in the severity of the symptoms induced by the different *Fusarium* species, with isolates of *F. oxysporum*, *F. avanaceum* and *F. redolens* inducing the most severe symptoms. The findings were similar to those of Ren (2003), who reported *F. oxysporum* and *F. avanaceum* as being highly virulent; however, the same authors found *F. equesiti* to be highly virulent, whilst our study concluded *F. equesiti* to be only moderately virulent, possibly due to variation in the faba bean genotypes used. This difference may also indicate the presence of different pathotypes of the same *Fusarium* species with different levels of virulence. This phenomenon is common in other fungal species, such as *Ascochyta rabiei*, which has four pathotypes with different levels of virulence (Reddy & Kabbabeh, 1985; Imtiaz et al., 2011). Accordingly, there is a need to determine how different faba bean genotypes react to different species of *Fusarium*, and how the same faba bean genotype reacts to different isolates from the same *Fusarium* species, to explore the presence of different pathotypes, which may have an important impact on disease management.

## Summary

In this study, we have identified the likely casual agents of foot and root rot in faba beans in England, and their relative pathogenicity, using a combination of molecular and microbiological approaches. The identification of dominant species and causal agents of the disease in the UK will help refine monitoring of the disease, enabling streamlined monitoring earlier in the growing season, and supporting the development of integrated disease management strategies. The variations in symptoms induced by the species in our study suggests that there could be genetic variation across faba bean varieties with implications for their resistance to different pathogen species. The sequences generated in our study could potentially be used to develop a soil risk prediction tool to identify the existence of different *Fusarium* species in the soil, in parallel with identification of resistant varieties to help farmers select the most appropriate variety to grow in infected locations.

## Acknowledgements

BA was funded by the Council for At-risk Academics (CARA) and the School of Natural and Environmental Sciences, Newcastle University, and thanks them for financial support throughout the project. We would like to thank the growers and producers who provided samples for this work in conjunction with the PGRO.

## Author contributions

Basem Attar: Conceptualization; Writing – original draft; Writing – review & editing; Investigation; Formal analysis; Visualization; Project administration. James J. N. Kitson: Conceptualization; Writing – review & editing; Investigation; Supervision; Project administration. Jordan P. Cuff: Writing – review & editing; Investigation; Formal analysis; Supervision; Visualization; Project administration. Becky Howard: Conceptualization; Methodology; Resources. Ana Lages: Methodology; Resources. Dina Gomez: Methodology; Resources. Neil Boonham: Conceptualization; Writing – review & editing; Investigation; Supervision; Funding acquisition; Visualization; Project administration.

## Data availability statement

All data and code are publicly available via Zenodo: https://doi.org/10.5281/zenodo.13952394 (Attar et al., 2024). All sequence data generated have been submitted to GenBank and are awaiting accession codes.

